# Evaluating the Therapeutic Efficacy of Iopanoic Acid in a DMM-Induced Osteoarthritis Mouse Model and Osteochondral Lesioned Human Explants

**DOI:** 10.1101/2025.07.15.663868

**Authors:** S. Sana Sayedipour, Giorgia Mazzini, Margo Tuerlings, Jelle Nikkels, Marijke Koedam, Luis J. Cruz, Rachid Mahdad, Louise de Weerd, Bram van der Eerden, Yolande FM Ramos, Ingrid Meulenbelt

## Abstract

**Objective:** To evaluate the therapeutic potential of iopanoic acid (IOP), a thyroid hormone pathway inhibitor, in preserving cartilage and bone integrity in osteoarthritis (OA), using *in vivo* and *ex vivo* tissue models.

**Design:** In the DMM mouse model, IOP was administered through intra-articular (i.a.) injection, either alone or combined with a thermosensitive hydrogel to enable sustained release. Histological analyses included Safranin O/Fast Green staining and OARSI scoring. Immunohistochemistry was performed for COL2, MMP13, and CCDC80 to evaluate anabolic, catabolic, and hypertrophic markers. Micro-CT assessed subchondral bone changes. In the *ex vivo* studies, IOP was applied to lesioned human osteochondral OA explants. Matrix degradation and repair were evaluated by sulfated glycosaminoglycan (sGAG) release, Mankin histology scores, and RT-qPCR for cartilage matrix genes.

**Results:** Administration of IOP significantly reduced cartilage degeneration in DMM mice (*P* ≤ 1.0×10^-4^), characterized by increased COL2, and decreased MMP13 and CCDC80 expression. Notably, IOP also prevented pathological subchondral bone thickening. In human explants, IOP treatment led to a significant reduction in sGAG release compared to untreated explants on day 6 of the IOP treatment. Moreover, Mankin scores were significantly improved in IOP-treated compared to untreated explants, indicating reduced cartilage degradation.

**Conclusion:** IOP demonstrates strong chondroprotective effects, reducing cartilage degradation and promoting repair in OA models. Its combination with a thermosensitive hydrogel amplifies therapeutic potential, offering a promising strategy for OA treatment. Next steps are to optimize delivery and validate early molecular effects.

## Introduction

Osteoarthritis (OA) is a prevalent, complex, disabling disease of articular joints, characterized by degradation of articular cartilage and remodeling of the subchondral bone [1]. Despite extensive research, no treatment has been found that can alter the course of the disease. Consequently, patients obtain palliative care until they are eligible to undergo an invasive joint replacement surgery at late stage disease. This has a significant impact on their quality of life and will further increase burden on society given the global aging population [2]. To develop innovative effective treatments, a deeper understanding of the underlying mechanisms of OA disease processes is essential [3, 4]. To gain such insights, genetic studies have been performed and have highlighted a significant role for genes involved in regulating chondrocyte transitions during endochondral ossification [4–6]. One such gene is *DIO2* encoding the enzyme deiodinase iodothyronine type 2 (D2) that converts intracellular inactive thyroxine (T4) into its active form, triodothyronine (T3), thereby increasing intracellular T3 bioavailability. Notably, T3 signaling is known to initiate terminal chondrocyte maturation in growth plate tissues hence inducing hypertrophy and leading to cartilage breakdown and mineralization [7, 8]. By applying functional follow-up studies of the *DIO2* gene in articular cartilage, it was shown that upregulation of *DIO2* expression [8, 9], affected articular cartilage integrity. Subsequently, *in vitro* [8] and *in vivo* [10] models were applied to demonstrate that attenuation of D2-enzyme activity by Iopanoic acid (IOP) could effectively prevent chondrocyte terminal maturation. Even more, we showed that IOP could have a beneficial effect to injurious mechanical stress inflicting OA-like damage in *ex vivo* aged human osteochondral explants [11]. To accommodate sensitive readout in such preclinical models we showed that CCDC80 encoding Coiled-Coil Domain Containing 80 is a robust and sensitive marker of T3 induced chondrocyte terminal maturation [12]. Together, we have previously shown that targeted inhibition of D2, could help preserve cartilage homeostasis by restraining chondrocyte hypertrophy and maintaining matrix integrity [8].

To facilitate localized and sustained delivery of IOP for therapeutic use, we previously developed and characterized a thermosensitive hydrogel based on 25% poloxamer 407 (P407), which enables minimally invasive intra-articular injection and drug retention at the joint site [13]. This hydrogel has proven to be effective for encapsulating small molecules like IOP, enhancing local bioavailability while minimizing systemic exposure. Building upon this, here we set out to evaluate the therapeutic potential of IOP in reducing OA-related damage using two complementary models: the *in vivo* destabilization of the medial meniscus (DMM) mouse model and *ex vivo* lesioned OA human osteochondral explants. The DMM model effectively replicates mechanical instability-induced OA progression in humans, providing a reliable platform for studying cartilage degradation and testing interventions [14]. In parallel, human OA explants retain pathophysiological features of diseased cartilage, enabling direct evaluation of therapeutic responses in clinically relevant tissues. This combined approach provides a robust framework to test IOP’s potential in modulating chondrocyte activity, preserving the cartilage extracellular matrix (ECM), and maintaining cartilage integrity in OA.

## Methods

### *In vivo* study design

30 male C57BL/6 J mice (11 weeks, 25 ± 2 g) were purchased from Charles River Laboratories (Charles River, Chatillon-sur-Chalaronne, France) for this study. The animal procedures were all conducted at the Leiden University Medical Center and were approved by the Animal Welfare Committee (IvD) under number AVD1160020171405 - PE.18.101.005. All mice were housed in groups in polypropylene cages on a 12-hour light/dark cycle with unrestricted access to standard mouse food and water A 7-day acclimatization period was provided prior to initiation of the experimental procedures.**Figure 1-A** represents the timeline of the *in vivo* mice experiment.

**Figure 1:**
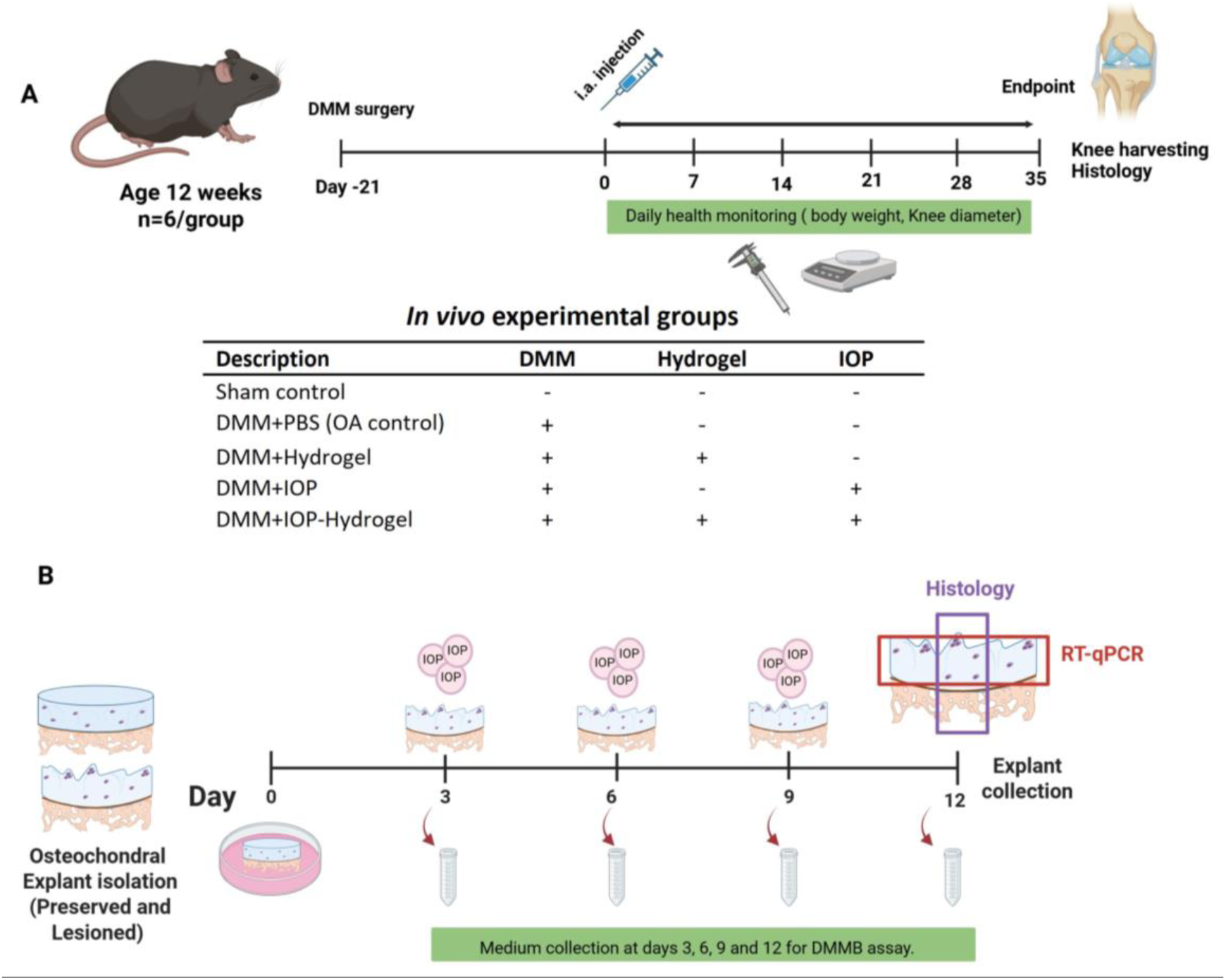
Experimental timeline: (A) *In vivo* OA model. 21 days before start treatment, we induce OA by cutting medial meniscus to develop OA. At day 0,treatment groups (PBS, Hydrogel, IOP+/- hydrogel) injected i.a. in to the mice knee. 35 days after injection, mice were sacrificed and their right knee collected to perform histology. (**B) *Ex vivo* experiment:** Macroscopically preserved and lesioned osteochondral explants were punched from the human OA knee and taken into culture. Lesioned osteochondral explants were treated with IOP (100μM) from day 3 onwards. Media was collected at the indicated days and subsequently each explant received fresh media. Finally, on day 12, explants harvested, one section was fixed in 4% formaldehyde for histology, while for the remaining explant cartilage and bone were separated, snap frozen and stored at -80°C for downstream analyses. DMM= Destabilization of the Medial Meniscus; i.a.= Intra articular; OA= Osteoarthritis; IOP= Iopanoic acid.

Surgical destabilization of the medial meniscus(DMM) was performed on the right knee joint in n = 24 male C57BL/6J mice at 12 weeks old, as described before [14, 15]. In brief, after isoflurane inhalation anesthesia in mice, a medial incision of the right knee joint was created to slightly shift the extensor muscle of the knee joint without transecting the patellar ligament and exposing the right knee joint. The MMLT was cut so that the medial meniscus could be moved to the medial side. After repositioning the knee extensor, the medial incision was sutured, and the skin closed. For mice that underwent sham surgery, a similar surgical approach was used. However, the knee joint was operated without MMTL transection. After surgery, mice had access to food and tap water. 21 days after DMM surgery, mice were randomly and equally divided into five groups (n = 6 in each group). They received one-time treatment via intra-articular (i.a.) injection, as described in **Supplementary Table S1**.

### Micro computed tomography measurements

35 days after i.a. administration of the treatment groups, mice were euthanized by continuous CO_2_ inhalation, the right knee joints were harvested, and fixed with 4% paraformaldehyde. After fixation for 48h, the specimens were transferred to 70% ethanol for high-resolution micro computed tomography (Micro-CT) (Skyscan 1072, Skyscan, Aartselaar, Belgium). The scanner was set at a gamma-ray voltage of 50 KV and a current of 200 uA, 0.5 mm Al filter, and a resolution of 9 μm per pixel. Subchondral bone morphology were analyzed using 3D data analysis software (CTAnalyzer, Skyscan). The region of interest (ROI) was defined within the subchondral bone of the medial tibial plateau and comprised 15 consecutive cross-sections, corresponding to a total ROI thickness of 135 µm. Quantitative parameters included bone volume fraction (BV/TV), trabecular thickness (Tb.Th), trabecular number (Tb.N), and trabecular separation (Tb.Sp) [16].

### Histological evaluation

For histological assessment, we followed the OARSI guidelines. The knee joints were fixed in 4% paraformaldehyde for 24h, followed by 5 days of decalcification in Mol-Decalcifier (Milestone) at 37°C. They were then embedded in paraffin, and knee joints were cut into 5μm sections. The sections were stained with Safranin O/Fast green and Hematoxylin & Eosin (H&E), and examined by Zeiss Axioscan Z1 slide scanner, using a 20x magnification to evaluate the cartilage damage of the femur and tibia in knee joint. The joint degeneration was assessed using the OARSI cartilage degeneration score in the histological assessment of the mice [14]. The scoring was performed by three independent researchers who were blinded to the conditions and to the scores of the other investigator. The results were averaged and the OA score of the sections was taken as the representative score of the knee joint, as described elsewhere [17, 18].

### Immunohistochemistry staining

Immunohistochemistry staining was performed on knee joint sections for collagen type 2 (COL2), MMP13 and CCDC80. For both types of staining, slides were blocked for endogenous peroxidase using 0.3% H_2_O_2_ for 10 min at room temperature. Antigen retrieval was performed with 25ug/ml Proteinase K (Prot K) prepared in 0.1M tris/HCL, pH 5.0 for 10min at 37°C, followed by 30min treatment with hyaluronidase (5 mg/ml in Tris/HCL pH 5.0). For both types of staining, all slides were blocked in 5% PBS-BSA for 30min at room temperature. Primary COL2 antibody incubation was done overnight at 4°C with 0.2µg/mL COL2 mouse monoclonal antibody (ab34712, Abcam, Cambridge, MA or with 0.2 µg/mL normal mouse IgG1 as isotype control (sc3877, Santa Cruz, Dallas, TX, USA). Primary MMP13 antibody incubation was done overnight at 4 C with 2 µg/mL MMP13 monoclonal antibody (SC-515284, Santa Cruz Biotechnology, Santa Cruz, Dallas, TX, USA) or with 2µg/mL normal mouse IgG1 as isotype control (SC-3877, Santa Cruz, Dallas, TX, USA). Primary CCDC80 antibody incubation was done overnight at 4 C with 2µg/mL CCDC80 polyclonal antibody (A65521, Thermo Fisher Scientific, Waltham, MA, USA) or with 2 µg/mL normal mouse IgG1 as isotype control (sc3877, Santa Cruz, Dallas, TX, USA). All antibodies were diluted in 5% PBS-BSA. The next day, slides were incubated with anti-mouse HRP (Envision, Dako, CA, USA) for 30min at room temperature and subsequently incubated with liquid DAB + 2-component system (Agilent, Santa Clara, CA, USA) for 5min. Sections were counterstained with hematoxylin, dehydrated, cleared in xylene and cover slipped with mounting medium (Sigma-Aldrich, Saint Louis, USA). Quantification of image intensity was performed using ImageJ, as described elsewhere. [19].

### *Ex vivo* human lesioned explant experiment

#### Study design and culture components

Osteochondral explants were collected from macroscopically preserved and lesioned areas of the femoral condyle of human OA knee joints within 3 hours of joint replacement surgery as described elsewhere [11]. A total of 38 osteochondral explants were obtained from six donors for this study and were divided into treatment groups as follows: control preserved (n=10 explants, N=6 donors), lesioned (n=11 explants, N=6 donors), and lesioned with IOP (n=16 explants, N=6 donors). Donor characteristics are provided in **Supplementary Table S2**. After harvesting, the explants were first washed with PBS and then maintained in a serum-free chondrogenic differentiation medium. This medium consisted of DMEM supplemented with ascorbic acid (50 μg/ml), L-proline (40 μg/ml), sodium pyruvate (100 μg/ml), dexamethasone (0.1 μM), ITS+, and antibiotics (100 U/ml penicillin and 100 μg/ml streptomycin) (all from Sigma-Aldrich, Zwijndrecht, The Netherlands) and was incubated at 37°C with 5% CO₂. The medium was refreshed every three days. IOP treatment for the lesioned explants began on day 3, with 100 µM of IOP added to the medium on days 3, 6, and 9, as illustrated in **Figure 1-B**. On day 12, explants were harvested for gene expression analysis, and the medium was collected for protein measurements.

### Determining cartilage integrity

#### Sulphated glycosaminoglycan (sGAGs) measurement

Sulphated glycosaminoglycans (sGAGs) concentration was measured in conditioned media of explants collected from day 3, 6, 9 and 12 following extraction with the photometric 1,9 dimethylene blue (DMMB; Sigma-Aldrich) dye method [20]. Shark chondroitin sulfate (Sigma-Aldrich) was used as the reference standard. To measure concentrations, 100 µl of medium or digested cartilage was mixed with 200 µl of DMMB solution and the absorbance at 525nm and 595nm was measured in a microplate reader (Synergy HT; BioTek, Winooski, USA).

#### Histology

Osteochondral explants were fixed in 4% formaldehyde for 48h and decalcified using Mol-Decalcifier (Milestone) for one weeks at 37°C. Subsequently, samples were dehydrated with an automated tissue processing apparatus and embedded in paraffin. Tissue sections of 5 μm were stained with Hematoxylin and Eosin (H&E), toluidine blue (Sigma-Aldrich), safranin-O/Fast Green, and mounted with Pertex (Sigma-Aldrich). Quantification of OA related cartilage damage was scored according to Mankin et al [21].

#### RNA isolation, Reverse Transcription and quantitative Real-Time PCR

Cartilage RNA was extracted by pulverizing the tissue using a Mixer mill 200 (Retch, Germany) and homogenizing in TRIzol reagent (Invitrogen, San Diego, CA). RNA was extracted with chloroform, precipitated with ethanol, and purified using the RNeasy Mini Kit (Qiagen, GmbH, Hilden, Germany). Genomic DNA was removed by DNase (Qiagen, GmbH, Hilden, Germany) digestion and quantity of the RNA was assessed using a Nanodrop spectrophotometer (Thermo Fischer Scientific Inc., Wilmington, USA). Synthesis of cDNA was performed using 200 ng of total mRNA with the First Strand cDNA Synthesis Kit (Roche Applied Science, Almere, The Netherlands) according to the manufacturer’s protocol. Subsequently, pre-amplification was performed and gene expression was determined with the QuantStudio 6 (Thermo Fisher) using Fast Start Sybr Green Master mix (Roche Applied Science). To measure changes in chondrocyte signaling upon perturbations, we measured gene expression levels by RT-qPCR. Primer sequences used are listed in **Supplementary Table 3**. Fold changes (FC) were determined using the 2-ΔΔCT method, in which cyclic threshold (CT) levels were adjusted for the housekeeping gene SDHA (-ΔCT) and subsequently for control samples (-ΔΔCT). A FC>1 represents an upregulation, while FC<1 depicts a downregulation.

#### Statistical analysis

For statistical analysis, GraphPad Prism 8.1.1 software (GraphPad Software, San Diego, CA, USA) was utilized. All data are expressed as the mean ± standard deviation (SD) of 3-5 independent repeated experiments, unless otherwise stated. Statistical significance was determined using Student’s t-test, unpaired, Mann-Whitney U test, and two-way analysis of variance (ANOVA). *P*-values were determined by applying a linear generalized estimating equation (GEE) to effectively adjust for dependencies among donors of the explants by adding a random effect for sample donor as we did not have perfect pairs for each analysisIn all analyses, a *P* -value ≤ a is considered an indicator of statistical significance and is expressed as: * *P* ≤ 0.05, ** *P* ≤ 0.01, *** *P* ≤ 0.001, **** *P* ≤ 0.0001.

## Results

### IOP slows-down cartilage degeneration in DMM mice model

H&E and safranin-O/Fast Green staining was applied to the knee sections of the outlined experimental groups (**Supplementary Table 1**) after performing the DMM model to assess OA related articular cartilage damage. As shown in **Figure 2A**, Safranin-O/Fast Green staining revealed preserved cartilage morphology in the sham group, although the red coloration indicating proteoglycan content appeared faint and less intense than typically expected. Despite this, the cartilage structure remained largely intact with smooth articular surfaces and clear tissue organization. Notable, the DMM+PBS group showed markedly disrupted cartilage architecture, with patchy and irregular safranin-O staining reflecting substantial proteoglycan loss and matrix degradation. These changes were accompanied by roughened cartilage surfaces and evident cracks, reflecting severe OA progression in the untreated PBS group. Treatment with hydrogel alone resulted in improved cartilage integrity compared to the PBS group, with smoother articular surfaces and more uniform matrix staining, although mild irregularities were still observed. The IOP-treated group showed similar improvements, with reduced structural damage. Notably, in the DMM+IOP-Hydrogel group cartilage surfaces appeared smooth and continuous, with reduced fibrillation and clearer tissue boundaries, suggesting a synergistic protective effect of IOP and the hydrogel delivery system. Quantitative scoring of meniscal damage (**Figure 2B**) confirmed that both IOP and hydrogel treatments significantly reduced OA-associated degeneration relative to PBS, as determined by unpaired t-tests (*P*≤1×10⁻^4^).

**Figure 2:**
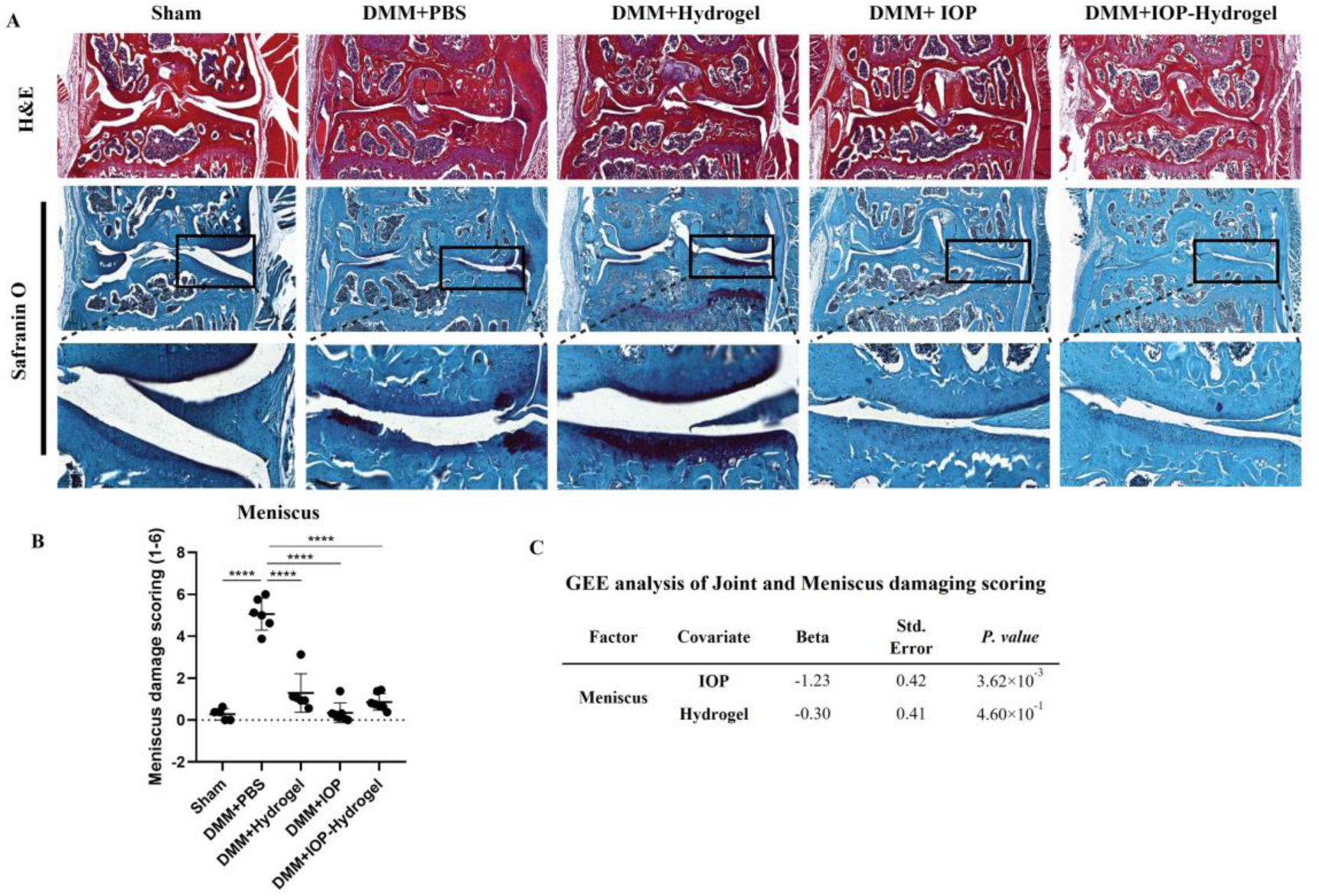
The evaluation of the therapeutic effect of the IOP and hydrogel on DMM mice model. Administration of experimental groups was performed 3 weeks after DMM surgery through one time i.a. injection. (**A**) Knee joints were harvested at 9 weeks after DMM surgery and analyzed histologically by H&E and Safranin O-fast green staining. Scale bars, 40μm. (**B**) Quantification of the meniscus lesions were graded on a scale of 0–6 using the damage scoring system. (**C**) Multivariate analyses of damage scoring using GEE to understand the independent therapeutic effect of IOP and hydrogel in the meniscus lesions. All data are shown as means ± standard deviations (n= 6). \*\*\*\**P*≤ 0.0001. DMM= Destabilization of the Medial Meniscus, H&E= Hematoxylin and Eosin; i.a.= Intra-articular; GEE= Generalized Estimating Equation.

To further dissect the independent contributions of IOP and hydrogel, a GEE analysis was performed (**Figure 2C**). The results revealed a significant protective effect of IOP on meniscal integrity (beta=– 1.23, *P*=3.62×10⁻^3^), while the hydrogel treatment alone did not reach statistical significance (beta=– 0.30, *P*=4.60×10⁻^1^). These findings highlight the chondroprotective efficacy of IOP and suggest that, although the hydrogel alone may offer mild structural support, its primary value lies in facilitating localized and sustained delivery of therapeutic agents such as IOP.

### IOP modulates COL2, MMP13, and CCDC80 expression in meniscus lesions of the DMM mice

To further investigate the therapeutic effects of IOP and/or hydrogel treatment on cartilage homeostasis, we examined markers of cartilage matrix synthesis, matrix degradation, and terminal chondrocyte maturation. Specifically, we performed immunohistochemical (IHC) staining for an anabolic extracellular matrix protein (COL2), a catabolic protease involved in matrix breakdown (MMP13), and a marker of terminal chondrocyte differentiation (CCDC80) (**Figure 3A**). COL2 staining was intense and continuous in the sham group but markedly reduced in the DMM+PBS group, indicating a significant loss of cartilage integrity. In contrast, MMP13 and CCDC80 expression were elevated in the DMM+PBS group, particularly in the deep cartilage zones and subchondral areas, reflecting increased catabolic activity and chondrocyte hypertrophy, respectively. Quantitative analysis (**Figure 3B**) confirmed these observations, showing a significant decrease in COL2 (*P* < 1×10⁻^4^) and significant increases in MMP13 and CCDC80 staining (*P* ≤ 1×10⁻^4^ for both) in the DMM+PBS group compared to sham.

**Figure 3:**
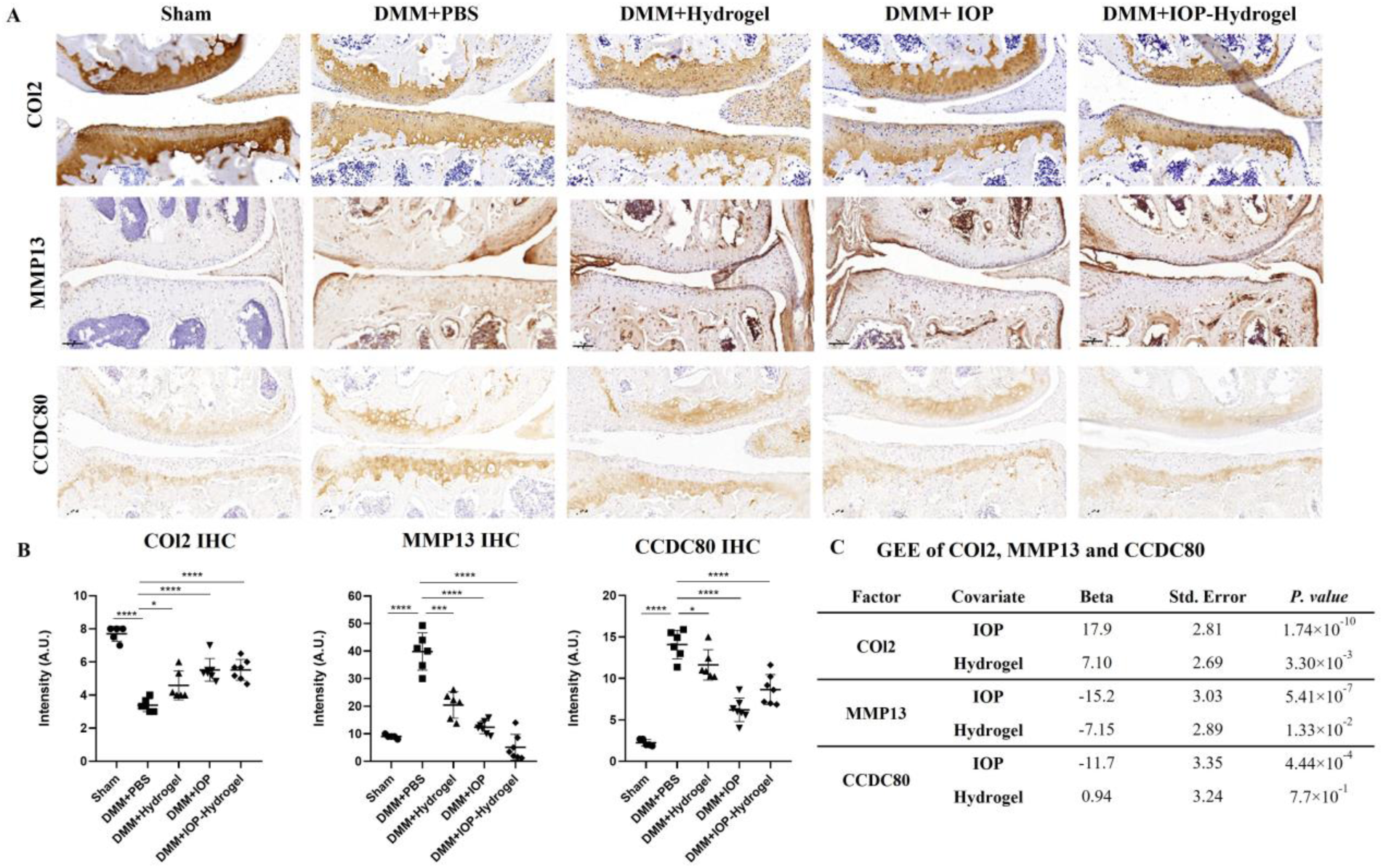
Immunohistochemistry staining of COl2, MMP13, and CCDC80 of the medial condyle cartilage at 9 weeks after the DMM procedure. (**A**) Representative IHC images of COl2, MMP13, and CCDC80 staining in meniscus lesions of the knee joint. (**B**) Quantification of IHC staining intensity for COl2, MMP13, and CCDC80. (**C**) Multivariate analyses of COl2, MMP13 and CCDC80 using GEE to understand the independent therapeutic effect of IOP and hydrogel in the meniscus lesions. All data are shown as means ± standard deviations (n= 6). \**P*≤0.05, \*\*\*\**P*≤0.0001. GEE: Generalized Estimating Equation.

GEE analysis (**Figure 3C**) confirmed that IOP treatment significantly increased COL2 staining intensity (beta =17.9, *P*=1.74×10⁻¹⁰), indicating a strong enhancement of cartilage matrix synthesis and structural integrity. Hydrogel treatment also led to a statistically significant increase in COL2 intensity (beta=7.10, *P* =3.30×10⁻³), though the effect was less pronounced than with IOP. For MMP13, both IOP and hydrogel treatments significantly reduced staining intensity. IOP showed a stronger suppressive effect (beta=–15.2, *P*=5.41×10⁻⁷) compared to hydrogel (beta=–7.15, *P*=1.33×10⁻²), indicating that both interventions inhibit catabolic activity in OA cartilage. With respect to CCDC80, IOP treatment significantly reduced its expression (beta=–11.7, *P* =4.44×10⁻⁴), suggesting that IOP can attenuate the progression of chondrocyte hypertrophy. In contrast, hydrogel treatment had no significant effect on CCDC80 staining (beta=0.94, *P*= 7.70×10⁻^1^).

### *In vivo* Micro-CT analysis revealed DMM-operated mice exposed to IOP exhibit reduced subchondral bone mass

Micro-CT analysis was conducted to quantitatively evaluate subchondral bone remodeling in the tibial plateau of mice. The region of interest (ROI) was defined in the subchondral trabecular bone, as illustrated in representative micro-CT images (**Figure 4A**), and analyzed using standard trabecular parameters. **Figure 4B** presents quantitative micro-CT data of trabecular parameters. Compared to the sham group, the DMM + PBS group exhibited notable deterioration in trabecular bone structure. Specifically, there was a significant increase in bone volume fraction (BV/TV%) (*P* = 2.28 × 10⁻²) and trabecular thickness (Tb.Th) (*P* = 2.81 × 10⁻²). Although trabecular number (Tb.N) showed a decrease and trabecular separation (Tb.Sp) also increased in the DMM + PBS group, these changes did not reach statistical significance. Treatment with IOP, either alone or in combination with hydrogel, normalized all measured trabecular parameters, with no statistically significant differences compared to the sham group. Notably, hydrogel-based treatments alone demonstrated additional effects. The DMM + hydrogel group showed a significant increase in Tb.N (*P* = 8.50 × 10⁻³) and a significant decrease in Tb.Sp (*P* = 3.20 × 10⁻³) compared to the DMM + PBS group, indicating partial restoration of the trabecular microarchitecture. Moreover, the DMM + IOP - hydrogel group exhibited a significant decrease in Tb.N (*P* = 1.31 × 10⁻²) and a significant increase in Tb.Sp (*P* = 1.30× 10⁻³) when compared to the DMM + hydrogel group, suggesting that IOP contributed to a normalizing effect, bringing these parameters closer to those observed in the sham group.

**Figure 4:**
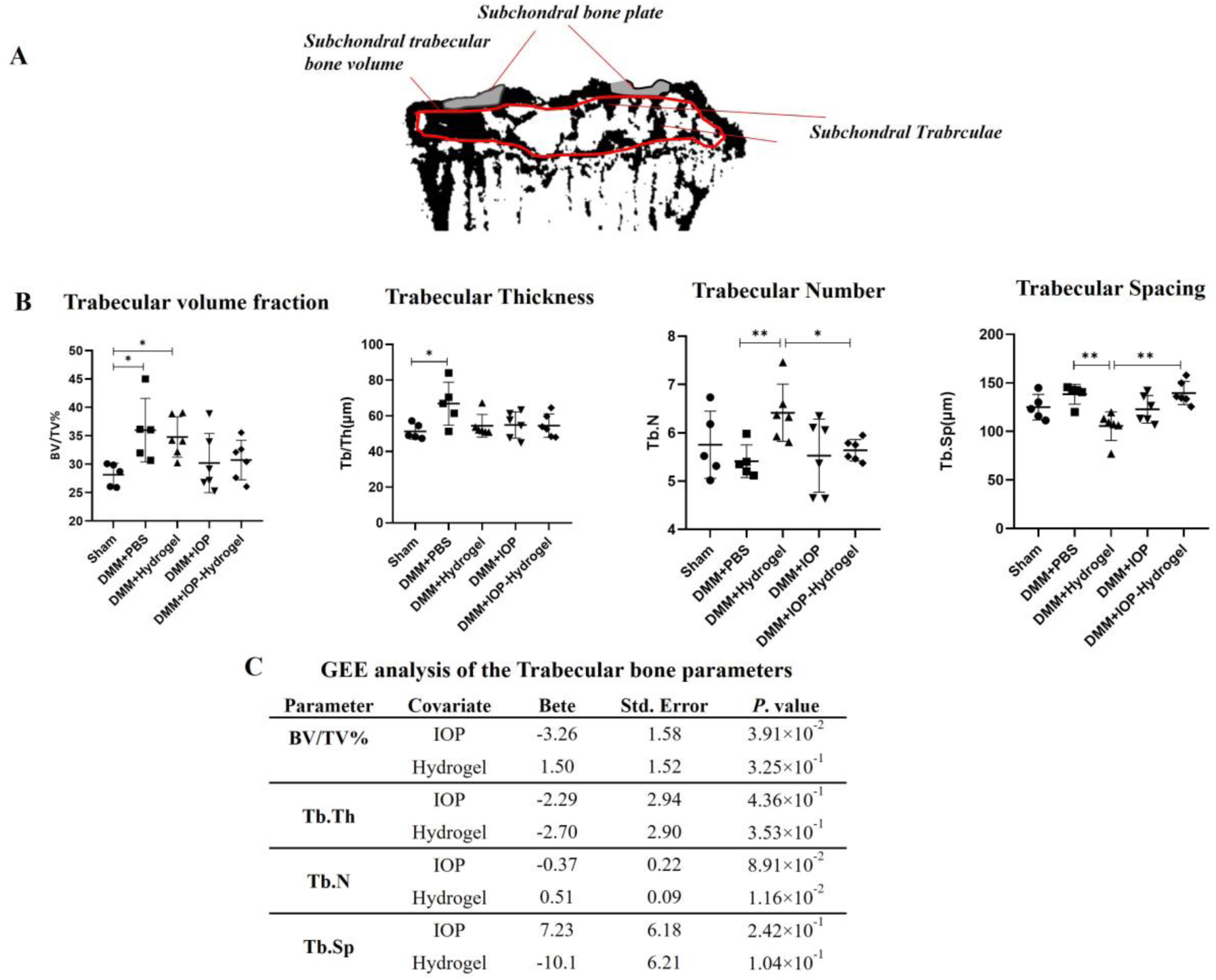
Micro-CT analysis of subchondral bone changes following IOP and hydrogel treatment in the DMM model. (**A)** Representative micro-CT images showing the defined region of interest (ROI) in the medial tibial plateau for trabecular bone analysis (highlighted in red). (**B**) Quantitative assessment of trabecular bone parameters, including bone volume fraction (BV/TV), trabecular thickness (Tb.Th), trabecular number (Tb.N), and trabecular spacing (Tb.Sp) across treatment groups. (**C**) GEE analysis indicating the effects of IOP and hydrogel on trabecular bone parameters. Statistically-significant differences are indicated by \**P*≤0.05, and \*\**P*≤0.01 between the indicated groups. GEE= Generalized Estimating Equations

To further evaluate the independent effects of hydrogel and IOP treatments, a GEE analysis was conducted (**Figure 4C**). Based on this analysis, both treatments appeared to normalize changes in subchondral bone parameters upon DMM-induced damage (**Figure 4C**). IOP treatment significantly reduced BV/TV% (Beta=–3.26; *P*=3.91×10⁻^2^), indicating strong suppression of OA-associated subchondral bone thickening. In contrast, hydrogel treatment significantly increased trabecular number (Tb.N) (Beta=0.51; *P* =1.16×10⁻²). Although most parameters did not reach statistical significance, this may be attributed to biological variability and relatively high variation. These findings suggest that IOP may exert a normalizing effect on trabecular bone architecture, particularly on BV/TV, while other changes may require larger sample sizes or extended treatment durations to achieve statistical significance, possibly due to smaller effect sizes and/or less robust outcomes.

### IOP enhances cartilage integrity of lesioned OA human osteochondral explants

To evaluate the translational potential of IOP treatment in human osteoarthritis, we applied IOP to lesioned osteochondral explants obtained from OA patients and compared its effects to both untreated lesioned explants and preserved (non-lesioned) biopsies. As a measure of cartilage matrix degradation, we quantified the release of sulfated glycosaminoglycans (sGAGs) into the culture medium on days 3, 6, 9, and 12 across all conditions (lesioned with/without IOP treatment and preserved explants). IOP treatment was initiated on day 3.

As shown in **Figure 5A**, lesioned explants released markedly higher levels of sGAGs compared to preserved explants, with the highest levels observed on day 3, indicating ongoing matrix degradation. Following IOP administration, sGAG release from the IOP-treated lesioned explants significantly declined by day 6, reaching levels comparable to preserved tissue. In contrast, untreated lesioned explants continued to exhibit elevated sGAG release. From day 6 onwards, sGAG levels gradually declined and stabilized in all groups.

**Figure 5:**
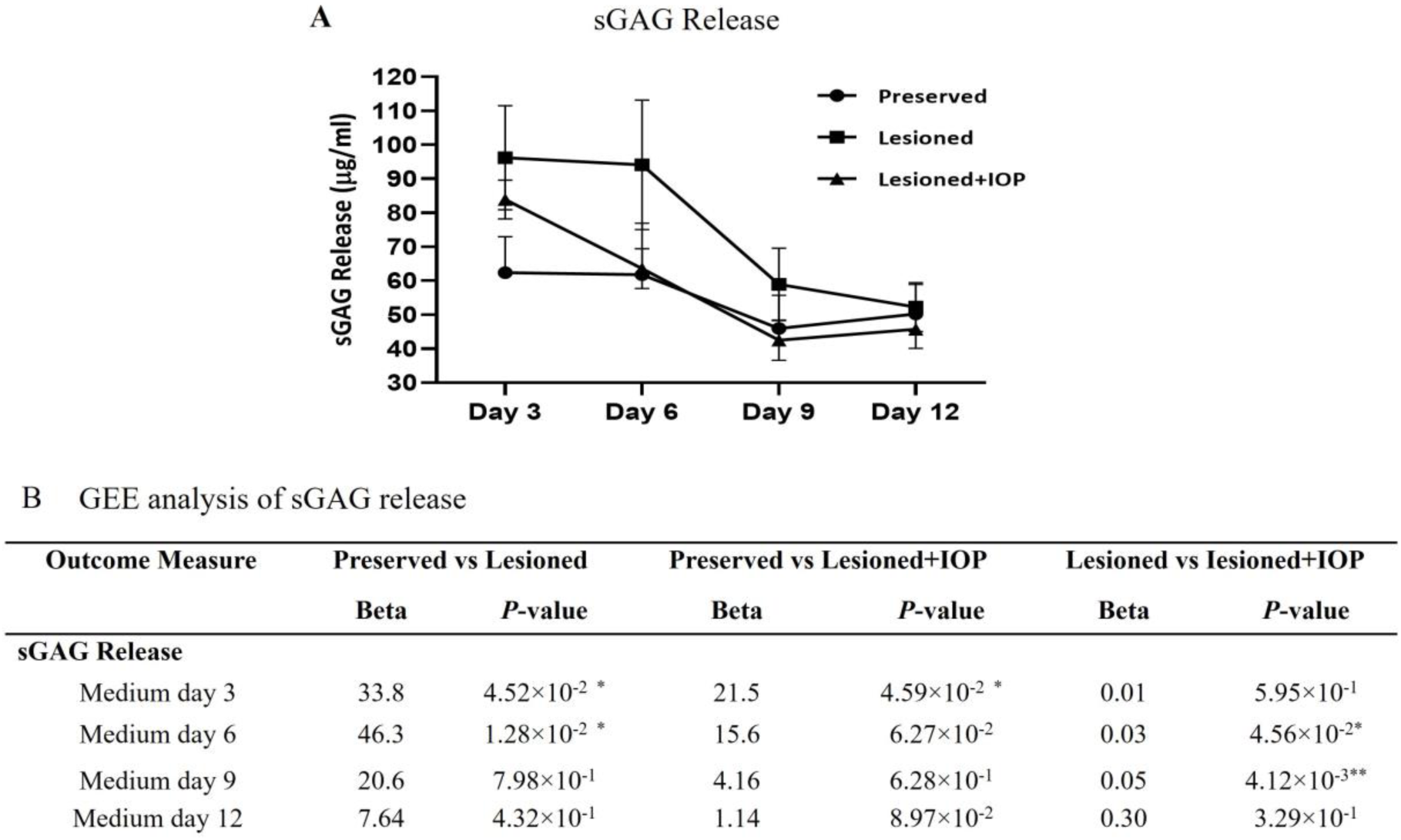
Assessment of sGAG release in preserved, lesioned, and IOP-treated lesioned human osteochondral explants. (**A**) sGAG release measured by the DMMB assay at days 3, 6, 9, and 12 from preserved, lesioned, and IOP-treated lesioned explants. (**B**) *P-*values of mean differences between controls and treated explants were estimated by GEE. Statistically-significant differences are indicated by *where *P*≤ 0.05, \*\**P*≤0.01 between the indicated groups. GEE= Generalized Estimating Equations.

GEE analysis (**Figure 5B**) confirmed all these observations with significant differences in sGAG release when comparing untreated lesioned and preserved explants on day 3 (Beta=33.8, *P*=4.52×10⁻²) and day 6 (Beta=46.2,*P* =1.28×10⁻²). IOP-treated lesioned explants also exhibited significant higher sGAG release on day 3 compared to preserved tissue (Beta=21.5, *P*=4.59×10⁻²), reflecting the pre-treatment state. Importantly, direct comparisons between untreated and IOP-treated lesioned explants revealed a significant therapeutic effect of IOP, with lower sGAG release at day 6 (Beta=0.03, *P*=4.56×10⁻²) and day 9 (Beta=0.05, *P*=4.12×10⁻³), indicating effective reduction of cartilage matrix degradation within 3 to 6 days post-treatment. By day 12, this difference was no longer significant, likely due to overall matrix stabilization in long-term culture with nutrient-rich medium (**Figure 5**). These findings collectively indicate that, while non-treated lesioned explants undergo significant cartilage degradation as evidenced by elevated sGAG release, IOP treatment successfully moderated this degradation and restored sGAG release to levels comparable to preserved explants.

Next, to assess the potential restorative effects of IOP on cartilage integrity in lesioned OA human osteochondral explants, Mankin scoring was employed to evaluate histological features indicative of cartilage health on day 12 of sample collection. As shown in **Figure 6A**, preserved explants displayed well-organized cartilage architecture with uniform and intense staining across H&E, Toluidine Blue, and Safranin O–Fast Green, representing intact extracellular matrix integrity. In contrast, non-treated lesioned explants showed substantial cartilage damage, characterized by disrupted tissue architecture, reduced proteoglycan staining, and surface fibrillation, confirming severe matrix degradation. These structural changes were reflected by a significant increase in Mankin scores compared to preserved explants (*P* ≤ 1×10^-4^), confirming the severity of matrix degradation in the lesioned group (**Figure 6B**). Importantly, lesioned explants treated with IOP demonstrated partial restoration of cartilage integrity, as evidenced by improved Safranin O and Toluidine Blue staining, suggesting enhanced proteoglycan retention and reduced cartilage breakdown. Consistent with these observations, Mankin scoring revealed a significant reduction in cartilage damage in IOP-treated lesioned explants compared to untreated lesioned samples (*P* = 4.70×10^-3^), further supporting the protective and restoring role of IOP in human osteochondral tissue.

**Figure 6:**
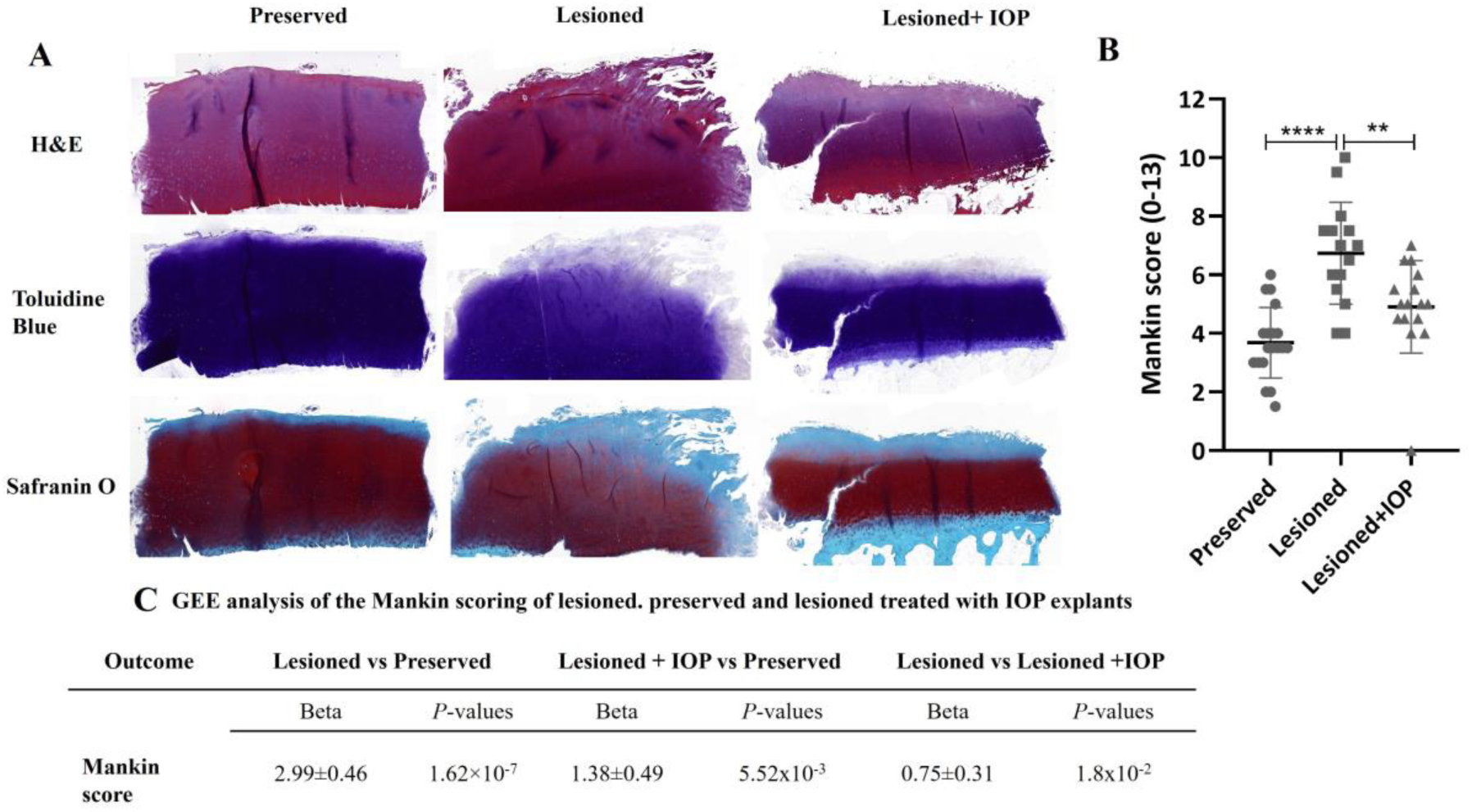
Histological evaluation and Mankin scoring of preserved, lesioned, and IOP-treated human lesioned osteochondral explants. (**A**) Representative histological images of H&E, Toluidine Blue, and Safranin O -fast green staining. (**B**) Quantification of cartilage degradation using the Mankin scoring system. Data are presented as scatter dot plots, with mean and 95% CI, and each circle represents a sample. (**C**) *P*-values of mean differences between controls and treated explants were estimated by GEE with Mankin score as dependent variable, treatment as factor and robust variance estimators to account for donor effects. \*\**P*≤0.01, \*\*\*\**P*≤0.0001. H&E: haematoxylin and eosin; GEE: Generalized Estimating Equation.

GEE analysis confirmed this observation, with lesioned explants displaying on average more damage than preserved ones (Beta = 2.99 ± 0.46, *P*=1.62×10⁻⁷). IOP-treated lesioned explants also exhibited higher Mankin scores than preserved explants (Beta=1.38±0.49, *P*=5.52×10⁻³), although the magnitude of degeneration was notably reduced. Importantly, direct comparison between lesioned and IOP-treated lesioned explants revealed a statistically significant reduction in cartilage damage following IOP treatment (Beta=0.75±0.31, *P*=1.8 × 10⁻²), confirming the therapeutic effect of IOP in reducing structural degeneration in human OA cartilage.

To further evaluate the effects of IOP treatment on chondrocyte phenotype, we performed RT-qPCR at day 12 (the final timepoint of the culture period) on chondral compartment of human osteochondral explants. A panel of genes associated with cartilage matrix synthesis and degradation (listed in **Supplementary Table S3**) was analyzed. *COL2A1* and *ACAN* were included as anabolic markers of cartilage matrix production, while *MMP13* was selected as a representative catabolic enzyme implicated in OA-related cartilage breakdown. Consistent with stabilization of the sGAG release and Mankin scoring data, no statistically significant differences in gene expression were observed among the preserved, lesioned, and IOP-treated lesioned groups at day 12 (**Figure 7**).

**Figure 7:**
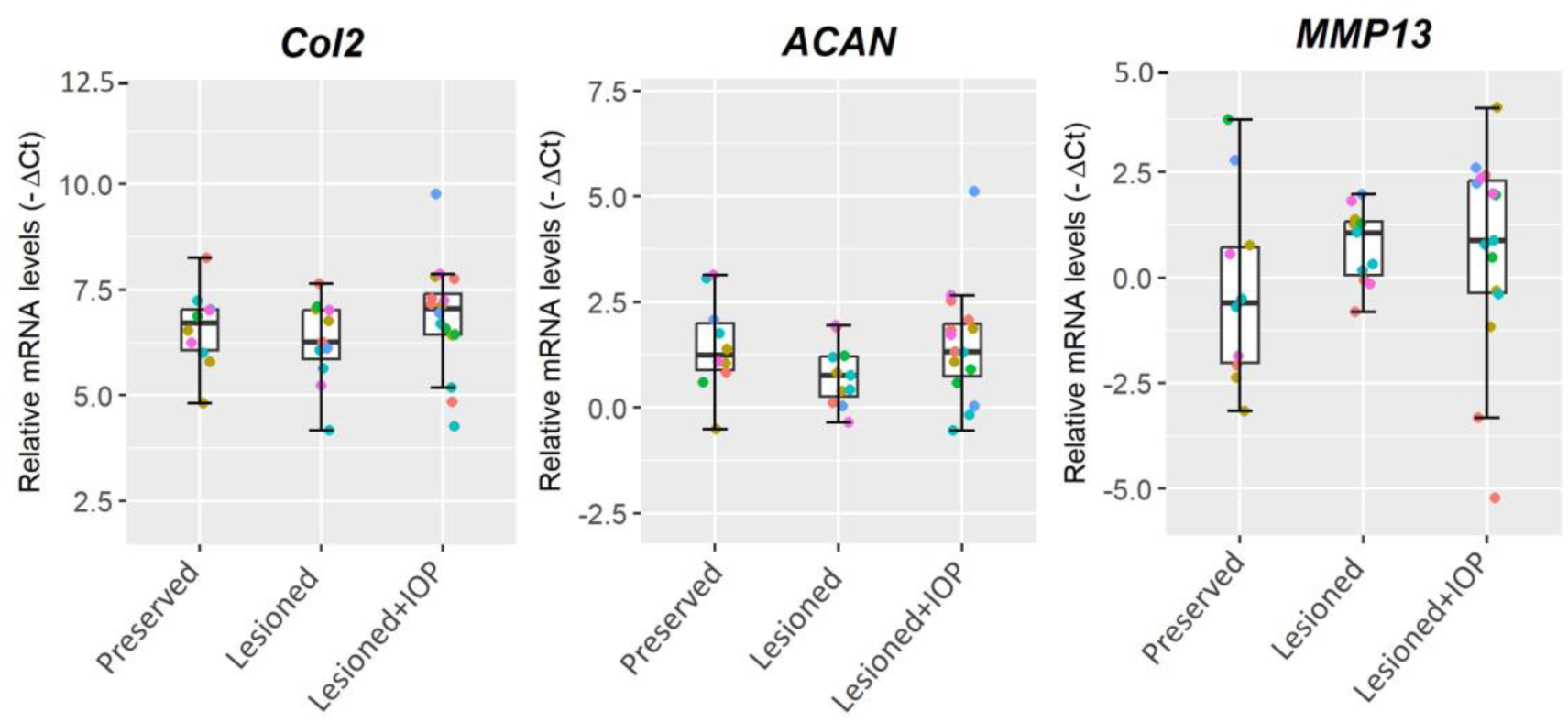
Relative mRNA expression levels of selected genes in preserved, lesioned, and IOP-treated lesioned human osteochondral explants after 12 days of culture using RT-qPCR. Abbreviations: see main text.

## Discussion

This study aimed to investigate the therapeutic potential of inhibiting D2 activity using the anti-deiodinase agent, IOP, in both an *in vivo* DMM-induced OA mouse model and a previously established *ex vivo* human osteochondral lesioned explant model. Our findings highlight that targeting D2 activity represents a promising strategy for slowing OA progression and may serve as an effective early treatment option. By limiting local thyroid hormone activation through D2 inhibition, IOP helps to reduce cartilage degradation and support tissue preservation, reinforcing the relevance of this pathway in OA pathology.

The DMM model mimics key aspects of human OA pathophysiology, particularly mechanical instability-induced joint stress and progressive cartilage degradation [22]. By utilizing this model, we were able to observe the disease progression and evaluate the impact of D2 inhibition in a controlled *in vivo* environment for the first time. Damage OARSI scoring demonstrated that IOP effectively reduced cartilage degradation in the DMM model. Additionally, the protective effects were more pronounced when IOP delivery was combined with a thermosensitive hydrogel. This effect likely stems from the hydrogel’s capacity to localize and sustain IOP delivery at the site of intra-articular injection. The hydrogel alone also showed mild protective effects, consistent with our previous findings. [23]

Immunohistochemical analysis further revealed that IOP COL2 expression, while reducing MMP13 and CCDC80 levels in cartilage. CCDC80 protein has emerged as a key player in the pathophysiology of OA, particularly in the context of chondrocyte hypertrophy and ECM dynamics. *CCDC80* gene expression significantly upregulated in hypertrophic chondrocytes, where it influences skeletal development and lipid metabolism, and is closely associated with the altered ECM remodeling characteristic of OA [12]. Importantly, recent findings highlight that the expression of *CCDC80* is markedly induced by T3, which accelerates chondrocyte hypertrophy and terminal maturation. These processes contribute to cartilage degradation, establishing CCDC80 protein as a robust and sensitive marker for hypertrophy in OA [12, 24, 25]. T3 signaling drives the terminal maturation of growth plate chondrocytes and primes cartilage to an OA-like disease state when exposed to environmental challenges such as mechanical loading [26]. These molecular changes suggest that IOP treatment not only protects cartilage from structural deterioration but also reshapes the chondrocyte phenotype toward a more regenerative, anti-hypertrophic state. Our findings are further supported by studies on aged human cartilage explants, where inhibition of thyroid hormone activation using IOP was shown to reduce the damaging effects of mechanical injury. These results suggest that IOP plays a protective role in maintaining cartilage homeostasis under mechanical stress [11].

In addition to assessment of cartilage integrity, we performed micro-CT analysis of mouse knee joints to evaluate the impact of IOP treatment on subchondral bone remodeling. Our findings demonstrate that, beyond its effects on cartilage, IOP also modulates the underlying bone architecture. Specifically, IOP treatment led to a significant reduction in bone volume, effectively counteracting the subchondral bone thickening typically associated with osteoarthritis progression. This aligns with the pathological features of OA, where early-stage disease is characterized by bone loss and cartilage degeneration, followed by excessive subchondral bone formation and osteophyte development in later stages [27]. However, when evaluating detailed trabecular parameters, the results were more complex. IOP alone was associated with increased trabecular spacing and decreased trabecular number, which may indicate impaired trabecular organization and reduced bone connectivity. These trends, while biologically relevant, were not statistically significant and were accompanied by considerable variability and relatively high standard errors. This variation may reflect differences in individual disease severity, response to treatment, or early-stage remodeling activity that is difficult to capture precisely in small cohorts. Hydrogel treatment alone showed moderately beneficial effects on trabecular microstructure. Importantly, the combination of IOP with hydrogel delivery appeared to reduce the adverse trends observed with IOP alone and promoted a more stable trabecular architecture. Taken together, these results indicate that while IOP can influence subchondral bone remodeling, the method of delivery plays an important role in shaping its overall effect, with hydrogel-based delivery offering a more balanced therapeutic outcome.

Building on the *in vivo* findings demonstrating IOP’s effectiveness in reversing OA pathology in the DMM mouse model, we further investigated its therapeutic properties using an *ex vivo* human osteochondral explant model from OA patients. Our findings confirmed that IOP treatment reduced cartilage degradation and promoted structural preservation in human OA lesioned explants. Biochemical analysis of sGAG release, quantified over 12 days via DMMB assay, confirmed these protective effects. By day 6 (3 days after starting with IOP treatment), sGAG release in IOP-treated lesioned explants were not significantly different from preserved controls, suggesting early stabilization of the ECM. This effect persisted through day 12, with treated explants maintaining sGAG release levels comparable to preserved tissue.

Histological evaluation, supported by Mankin scoring, further demonstrated IOP’s structural benefits. IOP-treated lesioned explants exhibited improved tissue organization, and increased proteoglycan retention. Although complete restoration to the morphology of preserved cartilage was not achieved, the significant reduction in Mankin scores in IOP-treated compared to untreated lesioned explants highlights IOP’s ability to reduce lesion-induced cartilage damage.

Despite the matrix-level improvements observed, RT-qPCR analysis at day 12 revealed no significant differences in the expression of key anabolic (*COL2A1*, *ACAN*) or catabolic (*MMP13*) markers across treatment groups. This lack of a detectable transcriptional response may be attributed to the timing of sampling, potentially missing early, transient gene expression changes. Additionally, the prolonged culture period in nutrient-rich medium may have facilitated partial recovery of the lesioned cartilage, further masking treatment-specific effects. This interpretation is supported by our sGAG release data, which showed that the most pronounced effect of IOP occurred within the first three days following treatment initiation. Therefore, it is likely that any early molecular regulatory effects of IOP had dropped by day 12. Future studies should include earlier timepoints, such as days 3 and 6 post-treatment, to more accurately capture the dynamic gene expression responses induced by IOP.

In summary, our findings from both *in vivo* mouse and human *ex vivo* models demonstrate that IOP treatment reduces cartilage degradation and supports tissue preservation in OA. In the DMM mouse model, IOP reduced structural damage, suppressed catabolic and hypertrophic markers, and promoted matrix integrity. Similarly, IOP preserved proteoglycan content in the human osteochondral explant model, reduced sGAG release, and improved cartilage histology in lesioned OA human explants. The early structural and biochemical improvements observed in both systems underscore the therapeutic potential of DIO2 inhibition and highlight the need for further investigation into the early molecular mechanisms of IOP in OA treatment.

## Acknowledgments

We thank all the participants of the RAAK study. The LUMC has and is supporting the RAAK study. We thank all the members of our group for valuable discussion and feedback. We also thank Enrike van der Linden, Demiën Broekhuis, Peter van Schie, Shaho Hasan, Maartje Meijer, Daisy Latijnhouwers, Anika Rabelink-Hoogenstraaten, and Geert Spierenburg for collecting the RAAK material. Furthermore, we thank all participating patients and doctors at Alrijne Leiderdorp.

## Author Contributions

Sana S. Sayedipour: Collection and/or assembly of data (*in vivo* and *ex vivo* experiments), data analysis and interpretation, manuscript writing, final approval of manuscript.

Giorgia Mazzini: Collection and/or assembly of data (*ex vivo* experiments), final approval of manuscript.

Margo Tuerlings: Collection and/or assembly of data (*ex vivo* experiments), final approval of manuscript.

Jelle Nikkels: Collection and/or assembly of data (*in vivo* experiments), final approval of manuscript. Marijke Koedam: Collection and/or assembly of data, final approval of manuscript.

Luis J. Cruz: Conception and design, final approval of manuscript. Rachid Mahdad: Conception and design, final approval of manuscript. Louise de Weerd: Conception and design, final approval of manuscript.

Bram van der Eerden: Collection and/or assembly of data, final approval of manuscript. Yolande FM Ramos: Conception and design, final approval of manuscript.

Ingrid Meulenbelt: Conception and design, data analysis and interpretation, manuscript writing, final approval of manuscript.

## Supplementary material

**Supplementary Table S1.** *In vivo* experimental groups

| Description | DMM | Hydrogel | IOP | Number |
| --- | --- | --- | --- | --- |
| Sham control | - | - | - | 6 |
| DMM+PBS (Control OA) | + | - | - | 6 |
| DMM+Hydrogel | + | + | - | 6 |
| DMM+IOP | + | - | + | 6 |
| DMM+IOP-Hydrogel | + | + | + | 6 |

DMM=Destabilization of medial meniscus, IOP= Iopanoic acid

**Supplementary Table S2.** Baseline information of the donors included in the *ex vivo* experiment this study.

| Characteristic | Average±SD [Range] |
| --- | --- |
| Age | 75.83±10.57 [57-87] |
| Sex | 3M, 3F |
| BMI | 30.83±3.067 [28-36] |

The table represents the age, sex, and BMI of donors used in this study. Legend: F=Female; age given in years)

**Supplementary Table S3.**
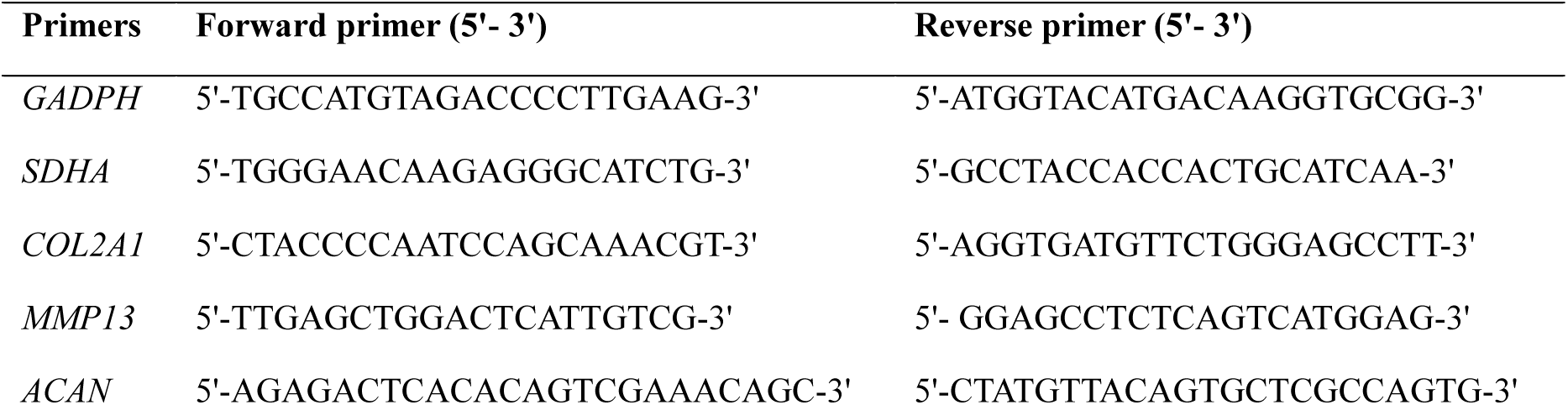
Primer sequences used in RT-qPCR

